# Culturomics of Aerobic Anoxygenic Phototrophic Bacteria in Wheat Phyllosphere Revealed a Divergent Evolutionary Pattern of Photosynthesis Genes in *Methylobacterium* spp

**DOI:** 10.1101/488148

**Authors:** Athanasios Zervas, Yonghui Zeng, Anne Mette Madsen, Lars H. Hansen

## Abstract

The phyllosphere is a habitat to a variety of viruses, bacteria, fungi and other microorganisms, which play a fundamental role in maintaining the health of plants and mediating the interaction between plants and ambient environments. A recent addition to this catalogue of microbial diversity was the aerobic anoxygenic phototrophs (AAPs), a group of widespread bacteria that absorb light through bacteriochlorophyll α (BChl *a*) to produce energy without fixing carbon or producing molecular oxygen. However, culture representatives of AAPs from phyllosphere and their genome information are lacking, limiting our capability to assess their potential ecological roles in this unique niche. In this study, we investigated the presence of AAPs in the phyllosphere of a winter wheat (*Triticum aestivum* L.) in Denmark by employing bacterial colony based infrared imaging and MALDI-TOF mass spectrometry (MS) techniques. A total of ~4480 colonies were screened for the presence of cellular BChl *a*, resulting in 129 AAP isolates that were further clustered into 25 groups based on MALDI-TOF MS profiling, representatives of which were sequenced using the Illumina NextSeq and Oxford Nanopore MinION platforms. Twenty draft and five complete genomes of AAPs were assembled belonging in *Methylobacterium, Rhizobium, Roseomonas* and a novel strain in Methylocystaceae. We observed a diverging pattern in the evolutionary rates of photosynthesis genes among the highly homogenous AAP strains of *Methylobacterium* (Alphaproteobacteria), highlighting an ongoing genomic innovation at the gene cluster level.

## Introduction

Plant-microbe interactions both above (phyllosphere) and below (rhizosphere) ground are common in nature. Traditionally, these relationships are investigated in the rhizosphere, where conditions are relatively stable and nutrient availability is rather high (Vorholt 2012; Carvalho & Castillo 2018). Despite recent work (reviewed in e.g. Lindow & Brandl 2003; Vorholt 2012), the microbe-phyllosphere (which is comprised by the aerial parts – and especially the leaves – of plants) interactions remain relatively understudied compared to rhizosphere. Models suggest that the total leaf surface is greater than 1 billion km^2^ (Woodward & Lomas 2004), potentially colonized by up to 10^26^ bacterial cells (Lindow & Brandl 2003).

Although the phyllosphere environment seems to be a common environment for bacteria to thrive in, it may be “extreme” for a plethora of reasons. First, different plants exhibit different growth patterns and climate adaptations. On one hand, annual plants complete their life cycle in just one growth season, whereas perennial plants grow and shed leaves every year. This creates a discontinuous, ever-changing habitat (Vorholt 2012). At the same time, environmental changes also affect the phyllosphere and its inhabitants. Winds, rainfall, frost, and drought all play important roles in shaping the conditions encountered by microorganisms of the phyllosphere. However, the most significant environmental driver is sunlight. Temperature differences in the upper leaves, especially those that form the canopy of various habitats may range up to 50°C between day and night. Sunlight, though beneficial for photosynthetic organisms, is also comprised of UV-radiation, which is damaging to the DNA of both prokaryotes and eukaryotes (Remus-Emsermann et al. 2014). In addition to the abiotic factors that make the phyllosphere a much more hostile environment compared to the rhizosphere, its inhabitants also face – directly or indirectly – the influence of their plant host. Leaves are surrounded by a thin, laminar, waxy layer, the cuticle, which renders the leaf surface hydrophobic, thus removing the excess of water that is collected due to rainfall, dew, or respiration from the stomata. This leads to the retention of little water most usually in veins and other cavities of the cuticle. These micro-formations also protect the colonizers from the surrounding environment (Beatie 2002). Apart from niche competition and scarce water availability, phyllosphere colonizers also need to compete for nutrients, as the cuticle makes the surface virtually impermeable to nutrients deriving from diffusion from the cells of the plant host (Wilson & Lindow 1994a, 1994b). At the same time, they also need to protect themselves from potential invaders and a plethora of antimicrobial compounds of prokaryotic or eukaryotic origin (Vorholt 2012).

Albeit the harsh conditions they encounter, phyllosphere microorganism exhibit remarkable biodiversity, which, in certain cases, can be comparable to that of the human gut microbiome (Xu & Gordon 2003; Knief et al. 2012). Through recent genomics and metagenomics studies, several bacterial genera, such as *Methylobacterium*, have been shown to be ubiquitous in the phyllosphere and their role in nutrient recycling, plant growth promotion and protection has been evidenced previously (Beattie & Lindow 1999; Gourion et al. 2006; Atamna-Ismaeel, Omri Finkel, et al. 2012; Kwak et al. 2014). One of the most interesting, recent findings in phyllosphere microbiota was the presence of aerobic, anoxygenic phototrophic (AAP) bacteria and rhodopsin-harboring bacteria in a variety of land plants (Atamna-Ismaeel, Omri M. Finkel, et al. 2012; Atamna-Ismaeel, Omri Finkel, et al. 2012). Employing 454 metagenome sequencing and also targeted PCR approaches for the identification of the chlorophyllide reductase subunit Y gene (*bchY*) and reaction center subunit M gene (*pufM*) it was possible to identify more than 150 bacterial rhodopsins belonging in 5 Phyla and a rich AAP community for the first time in non-aquatic habitats. AAP bacteria are commonly found in aquatic environment ranging from the arctic to the tropics and from high salinity lakes to pristine high altitude lakes (Yurkov & Hughes 2017). They rely on organic carbon compounds to cover their nutritional requirements and can also utilize light to produce energy in the form of ATP, without fixing carbon and without producing oxygen. Having been found in a variety of extreme environments and having shown their potential to outgrow non-AAP bacteria under nutrient limiting conditions, their presence in the phyllosphere may not be surprising. However, their relative abundance, at least in the rice phyllosphere (*Oryza sativa*), was up to 3 times higher compared to what is commonly found in marine ecosystems (Atamna-Ismaeel, Omri Finkel, et al. 2012).

These early culture-independent metagenomics studies provided valuable information about the presence of AAP bacteria in the phyllosphere. However, no pure cultures of AAPs have been described so far from phyllosphere and thus detailed genome information is lacking, which prevents an in-depth understanding of the ecological roles and evolutionary trajectory of AAPs in this unique niche. To expand our genomics views of this ecologically important group of bacteria in phyllosphere, we designed this study with the following aims: i) to create the first collection of AAP isolates from the phyllosphere, ii) to characterize their photosynthesis gene clusters (PGCs) and compare their gene content and molecular evolution, and iii) to expand the database of complete genomes of AAP bacteria from non-aquatic environments. Employing culturomics (Lagier et al. 2018) which combines colony infrared (IR) imaging systems, MALDI-TOF mass spectrometry, Illumina and Oxford Nanopore sequencing technologies, we were able to provide 20 draft genomes that contain complete photosynthetic gene clusters and 5 complete genomes that contain plasmids, including a novel strain in the Methylocystaceae family. The further comparison of the evolutionary trajectories of their PGCs revealed a diverging pattern among the highly homogenous *Methylobacterium* strains, highlighting an ongoing genomic innovation at the gene cluster level.

## Materials and Methods

### Sample collection and isolation of AAP strains

A piece of intact and alive winter wheat (*Triticum aestivum* L.) with a height of ca. 60 cm was collected from a field in Roskilde, Denmark, on 6 June, 2018 and transported to the lab for bacterial isolation on the same day. Wheat leaves were cut off and cleaned with running sterile water to remove dust and other temporary deposits from the ambient environment. Microbiota were collected from 8 pieces of leaves by rinsing the leaves in 80 mL PBS solution (pH 7.4) in a Ø 150-mm petri dish and scraping the leave surface repeatedly with sterile swabs until the PBS solution slightly turned greenish due to the fall-off of leave cells. From the supernatant, 14 mL were plated on 1/5 strength R2A plates (resulted in a total of 137 agar plates) and incubated at room temperature with normal indoor light conditions for two weeks. AAP bacteria were identified using a custom screening chamber equipped with an infrared imaging system, described in detail previously (Zeng et al. 2014). In short, plates were placed in a chamber sealed from light and exposed to a source of green lights. AAP bacteria containing bacteriochlorophyll absorb the green light and emit radiation in the near IR spectrum, which was captured by a NIR sensitive CMOS camera. The image was processed and the glowing colonies indicated the presence of BChl *a*. The colonies that give positive signals were marked, restreaked onto 1/5 strength R2A plates and incubated at room temperature for 72h. Verification of the light absorbing capabilities of the isolated strains was performed using the same setup. Strains that passed the verification step were further restreaked onto 1/5 strength R2A plates to ensure pure colonies of single AAP strains.

### Selection of AAP strains and whole genome sequencing

The isolated AAP strains were grouped and de-replicated using a MALDI-TOF mass spectrometer (Microflex LT, Bruker Daltonics, Bremen, Germany) before performing whole genome sequencing to reduce the cost while maintain the biodiversity. Briefly, a toothpick was used to transfer a small amount of an AAP bacterial colony onto the target plate (MSP 96 polished steel, Bruker) that was evenly spread out and formed a thin layer of biomass on the steel plate. The sample was then overlaid with 70% formic acid and allowed for air dry before addition of 1 uL MALDI-MS matrix solution (α-cyano-4-hydroxycinnamic acid, Sigma-Aldrich). A bacterial standard with well-characterized peaks (Bruker Daltonics) was used to calibrate the instrument. The standard method “MBT_AutoX” was applied to obtain proteome profiles within the mass range of 2-20 k Daltons using the flexControl software (Bruker). The flexAnalysis software (Bruker) was used to smooth the data, subtract baseline and generate main spectra (MSP), followed by a hierarchical clustering analysis with the MALDI Biotyper Compass Explorer software, which produced a dendrogram as the output for visual inspection of similarities between samples. An empirical distance value of 50 was used as the cutoff for defining different groups on the dendrogram, corresponding to different strains/species.

From each group on the dendrogram, one isolate was chosen and restreaked on ½ R2A plates and incubated at room temperature for one week. Prior to DNA isolation, the plates were again tested for the presence of light absorbing pigments as previously described. Bacterial colonies were scraped from the plates using a loop and immersed in 400μL PBS solution, vortexed until the cells had separated, spun down in a table-top centrifuge at 10.000g· at 4°C, the supernatant was removed and the pellets were re-dissolved in milliQ H_2_O. High molecular weight DNA was extracted from each strain using a modified Masterpure DNA purification kit (Epicentre) by replacing the elution buffer with a 10mM Tris-HCl - 50mM NaCl (pH 7.5 - 8.0) solution at the final step. DNA concentration was quantified on a Qubit 2.0 (Life Technologies) using the Broad Range DNA Assay kit and DNA quality was measured on a Nanodrop 2000C (Thermo Scientific). Whole genome shotgun sequencing was performed on all AAP strains on the Illumina Nextseq platform in house using the 2×150bp chemistry. Selected strains were also sequenced on the MinION platform (Oxford Nanopore Technologies, UK) in house using the Rapid Barcoding Kit (SBQ-RBK004) and the FLO-MIN106 flow cell (R9.4.1) following the manufacturer’s instructional manuals.

### Genome assembly and analyses

The Illumina pair end reads were trimmed for quality and ambiguities using Cutadapt (Martin 2011) and assembled using SPAdes (Bankevich et al. 2012). The resulting assemblies were analyzed using dRep (Olm et al. 2017) to compare their average nucleotide identities in order to identify clusters of closely related taxa. The assemblies were imported in Geneious R.11.2.5 (Biomatters Ltd.) and photosynthetic gene clusters were annotated using the *Roseobacter litoralis* strain Och149 plasmid pRL0149 (CP002624) and the *Tardiphaga* sp. strain G40 proteorhodopsin gene (AU collection – data not published) as references under relaxed similarity criteria (40%). The suggested genes of probable photosynthetic gene clusters were analyzed using BLAST (megablast, blastn) to verify their annotations. For full genome annotations, the assemblies were uploaded on RAST (Aziz et al. 2008; Overbeek et al. 2014; Brettin et al. 2015). The predicted, translated protein coding genes were imported in CMG-Biotools (Vesth et al. 2013) for pairwise proteome comparisons using the native blastmatrix program.

Hybrid assemblies of pair-end Illumina reads and long Oxford Nanopore Technologies reads were performed using the Unicycler assembler (Wick et al. 2017) utilizing the bold assembly mode. The resulting complete genomes were imported in Geneious, where their circularity was confirmed by mapping-to-reference runs using the short and long reads from the respective hybrid assemblies and the native Geneious mapper under default settings in the Medium-Low sensitivity mode. Whole genome alignments were performed using Mauve (Darling et al. 2004) and its progressiveMauve algorithm with default settings as implemented in Geneious.

### Phylogenetic analyses

Identification of the 16S ribosomal DNA sequences was performed by prodigalrunner in CMG-Biotools. Datasets containing the different genes from the identified photosynthetic gene clusters were created in Geneious. The resulting sequences were aligned using MAFFT v.7.388 (Katoh et al. 2002) using the FFT-NS-I x1000 algorithm with default settings as implemented in Geneious. Phylogenetic trees were created using RAxML (Stamatakis 2006) employing the GTR-GAMMA nucleotide substitution model and the rapid bootstrapping and search for best scoring ML-tree with 100 bootstraps replicates and starting from a random tree. Analysis of the photosynthetic gene clusters was performed on 5 selected genes that were present in all isolates, namely *acsF, bchL, bchY, crtB, pufL* and *pufM*. Individual gene alignments and phylogenetic trees were performed as described above. A super-matrix (8.240 characters) comprised of the total 7 gene alignments was created using the built-in “Concatenate alignments” function in Geneious, and a phylogenetic tree was created as above. All trees were visually inspected for disagreements. Finally, a phylogenomic tree was constructed in RAxML employing the GAMMA BLOSUM62 substitution model and the rapid bootstrapping and search for best scoring ML-tree with 100 bootstraps replicates and starting from a random tree, using as input the core protein alignment consisting of 6.988 characters created with CheckM (Parks et al. 2015) and its lineage_wf algorithm with default settings. Substitution rates were calculated as distances from the root, using the legend provided by the phylogenetic trees.

## Results

### Identification and isolation of Aerobic Anoxygenic Phototrophic Bacteria from the phyllosphere

The 129 phyllosphere isolates that were tested positive for the presence of bacteriochlorophyll-α were placed in 25 groups according to their protein mass profiles based on MALDI-TOF MS. A representative from each group were sequenced (Illumina) and their genomes assembled (SPAdes). Information about their phylogeny, total genome size, and genome assembly statistics are presented in table 1.

**Table 1:**
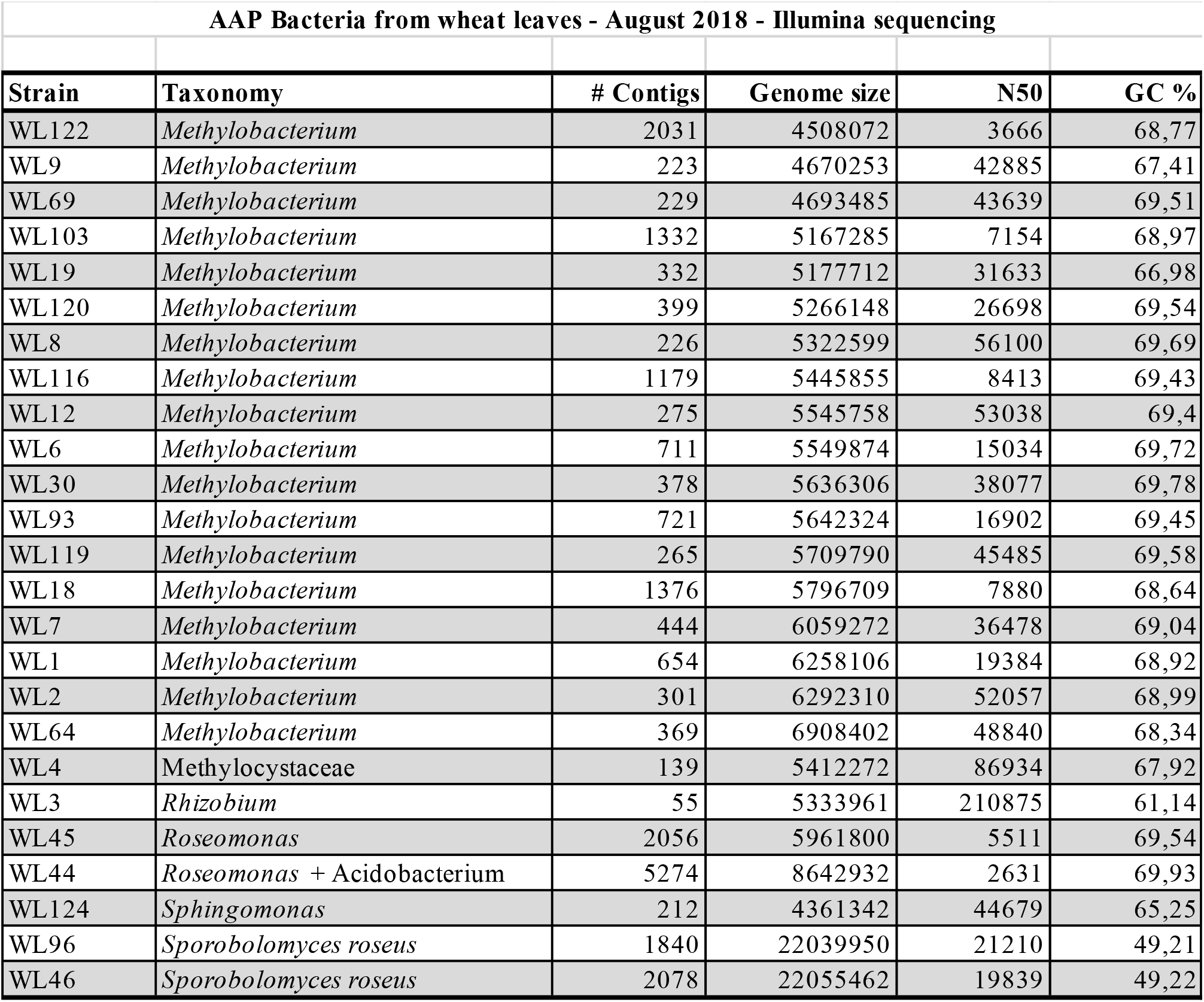
Genome statistics and phylogenetic information of the 25 isolates that were Illumina sequenced in this study

Analysis of the 16S region evidenced that isolate WL44 was in fact a mixture of 2 species (*Roseomonas* and an Actinobacterium), while isolates WL46 and WL96 actually were fungi (*Sporobolomyces roseus*). From the remaining 21 strains, 17 belonged in *Methylobacterium* (separated in 4 distinct groups), and the last 4 belonged in *Rhizobium, Roseomonas, Sphingomonas* and 1 novel species within the Methylocystaceae (WL4) (figure 1).

**Fig. 1:**
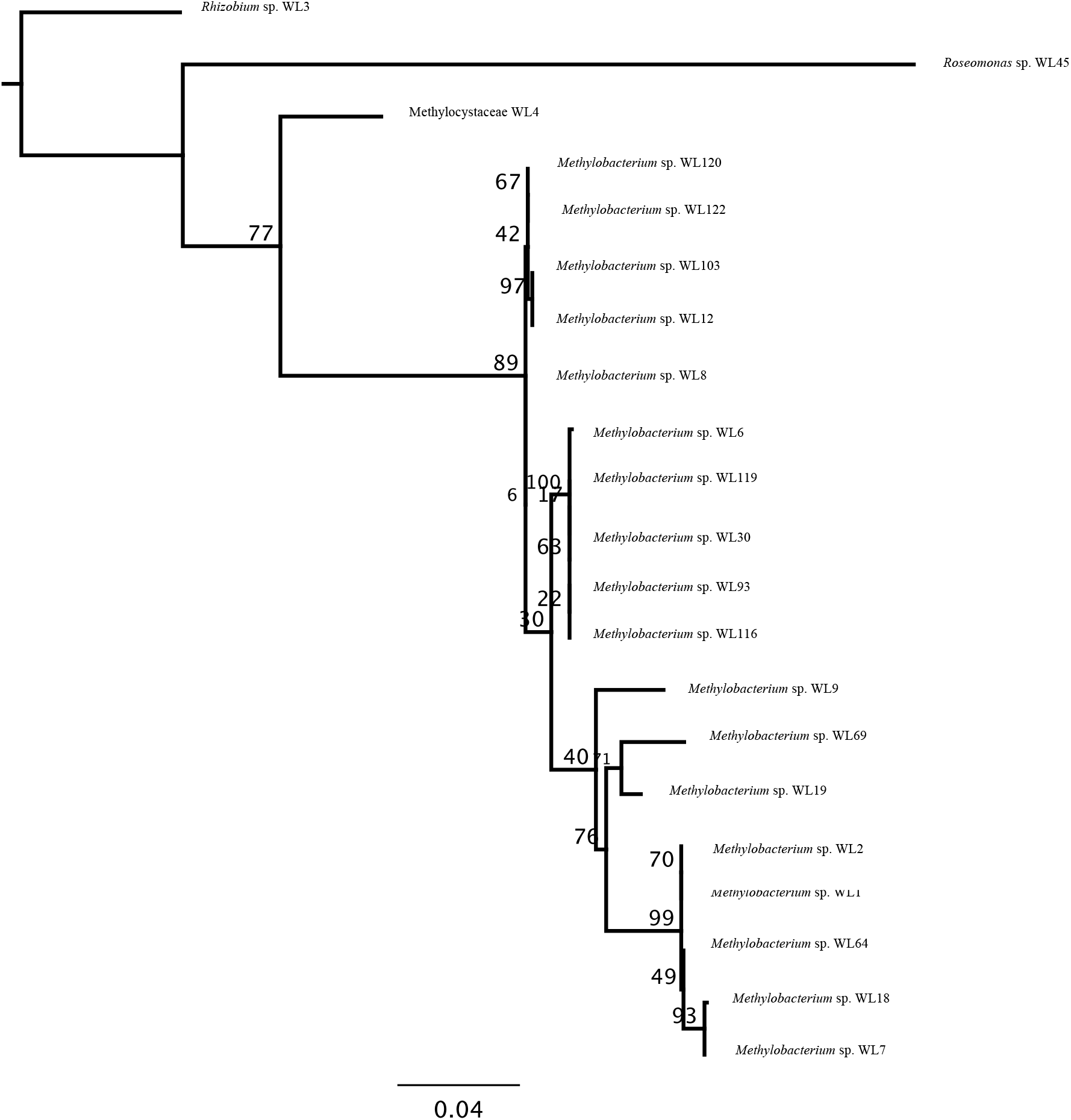
Phylogenetic tree based on the core proteome (CheckM) of the 21 pure bacterial isolates of the analysis. *Rhizobium* sp. WL3 was chosen as the outgroup. The legend shows substitution rates from the root of the tree and the node labels indicate bootstrap support values (%).

For WL4, analysis of the 16S region gave positive results on the family level, with most hits belonging to uncultured, environmental strains. The only characterized hits belonged to *Methylosinus trichosporium* (Gorlach et al. 1994) and *Alsobacter metallidurans*, a recently characterized novel genus, novel species (Bao et al. 2006). However, in both instances, the similarity was rather low (~98.5%). We included strains of both species in the core proteome analysis (CheckM – figure 2) and observed that WL4 is highly supported as sister to *Alsobacter* sp. SH9 (BS = 100%), both of which were distinct from both *Methylosinus trichosporium* OB3b and the remaining isolates, with very high bootstraps support values. The placement of WL4 was consistent, on this rather small dataset, throughout all the analyses.

**Fig. 2:**
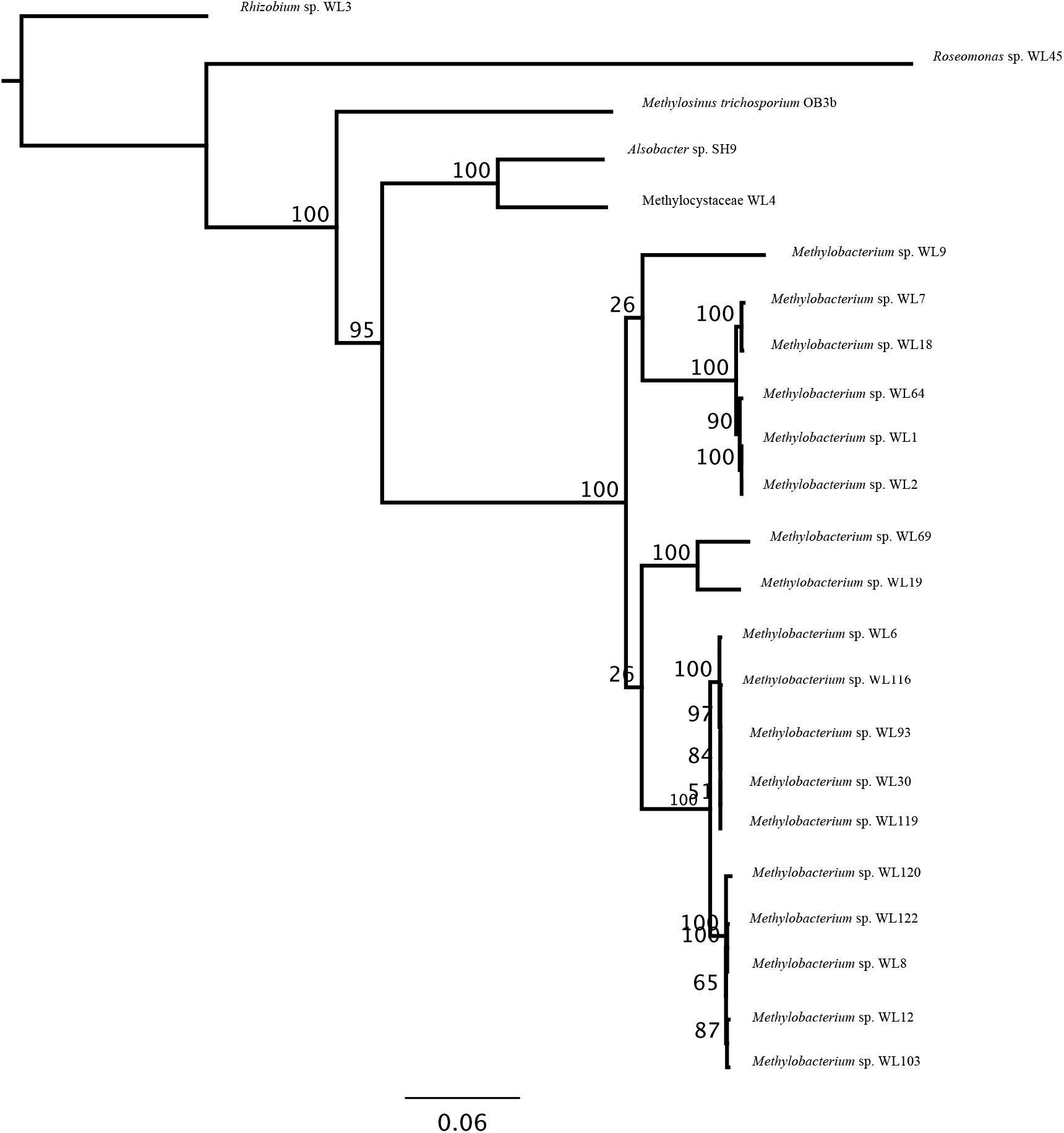
Phylogenomic tree based on the 16S region of the 21 pure bacterial isolates of the analysis. *Rhizobium* sp. WL3 was chosen as the outgroup. The legend shows substitution rates from the root of the tree and the node labels indicate bootstrap support values (%).

The 17 *Methylobacterium* isolates were further divided into 4 groups based on Average Nucleotide Identities (ANI) (figure 3). Group 1 is comprised of isolates WL9, WL19, and WL69, which probably comprise a distinct *Methylobacterium* species (ANI similarity < 85%). Group 2 is comprised of isolates WL1, WL2, WL64, WL7, and WL18, which belong to either the same species but different strains, or 2 different species (Species 2.1: WL1, WL2, WL64, Species 2.2: WL7, WL18). Group 3 is comprised of isolates WL6, WL30, WL93, WL116, WL119, and WL30, in 3 strains of the same species. Lastly, Group 4 is comprised of isolates WL8, WL12, WL103, WL122, and WL122, that are most likely 2 strains of the same species.

**Fig. 3:**
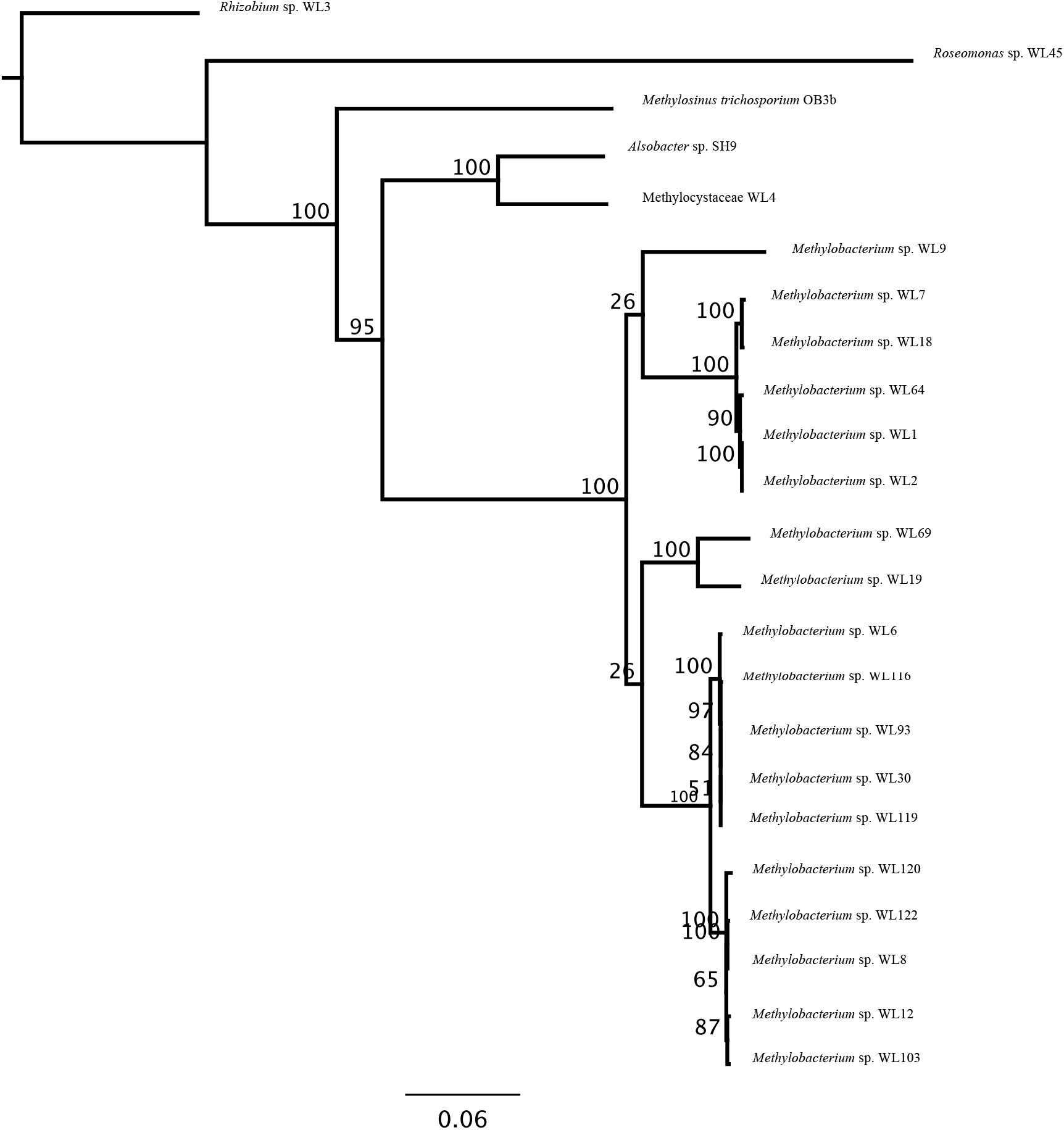
Whole genome, Average Nucleotide Identity (ANI) cladogram showing percentages of k-mer identities between the isolates.

This grouping is further supported by pairwise whole proteome comparisons (figure 4). For Groups 2 and 3, the average core protein content is ~ 4.500 proteins, while for Group 4 it is closer to 4.000. This may signify the presence of either more or larger plasmids that characterize the strains of this group. For Group 1, this number falls even more as in this case there clearly are 3 distinct species that make up the group.

**Fig. 4:**
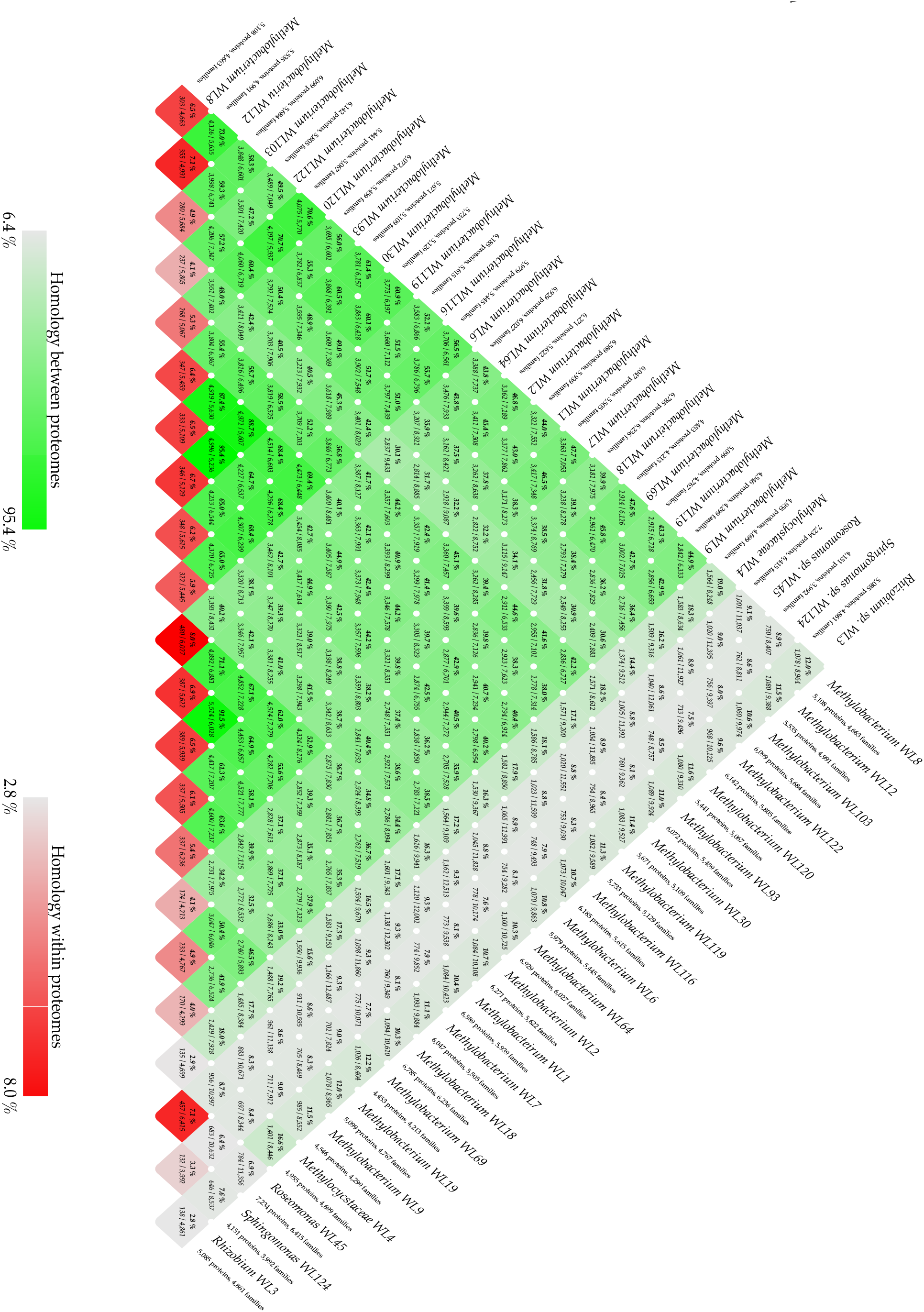
Pairwise proteome comparisons (BLAST-matrix) of the 21 bacterial isolates of the analysis. The grey-to-green color gradient indicates low to high percentages of identity. The bottom boxes show the presence of homologs within the same proteome (grey-to-red color gradient shows low to high presence of homologs in the specific proteome.

### Molecular evolution of the photosynthetic gene clusters

The photosynthetic gene clusters identified in the sequenced isolates all belong to RC-II type, meaning they contain bacteriochlorophyll-α and possess pheophytin – quinone type reaction centers (Xiong & Bauer 2002). All strains possess a full bacteriochlorophyll synthase gene set, except for strain WL4 that is lacking the *bchM* gene (table 2).

**Table 2:**
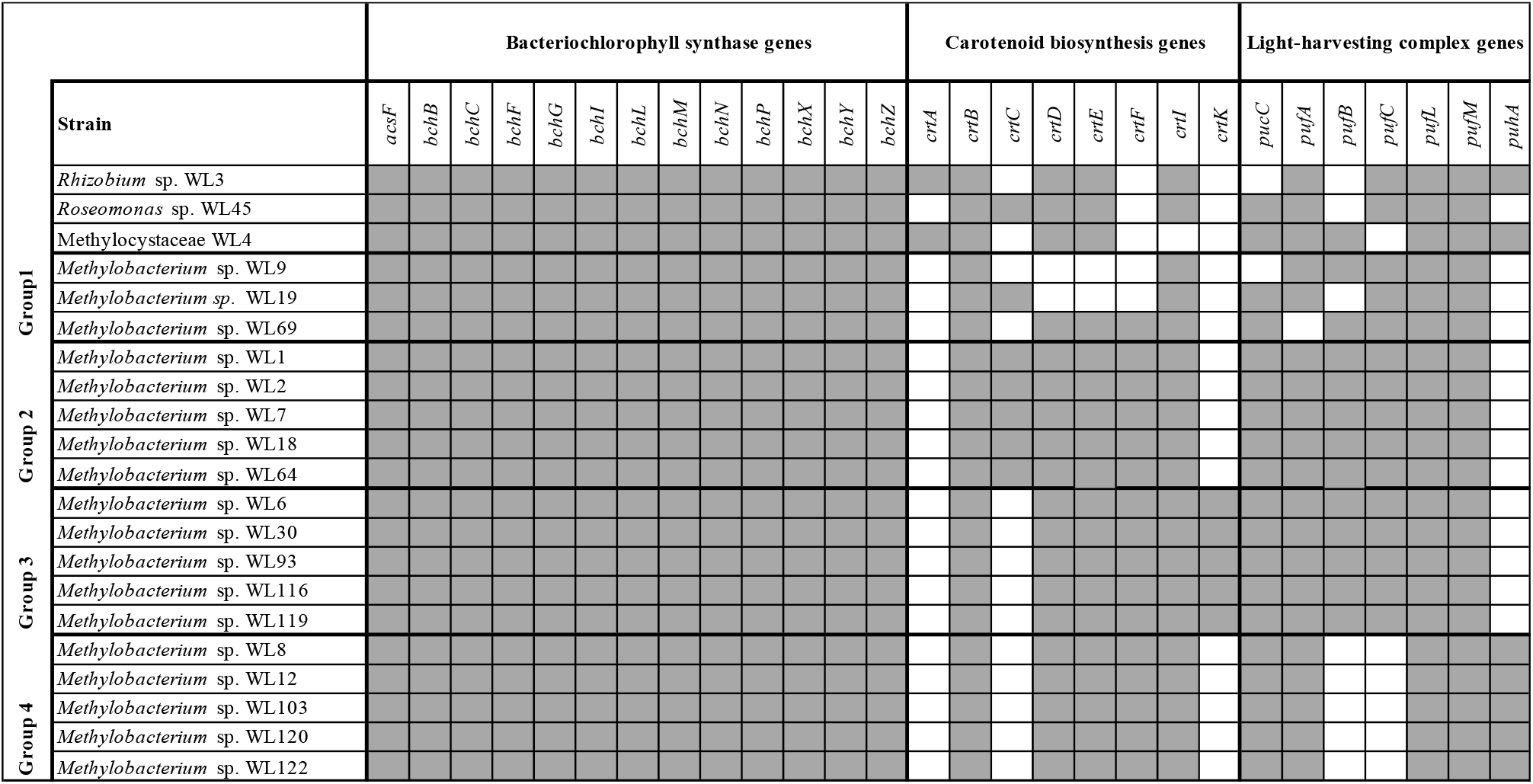
Gene content of the photosynthetic gene clusters identified in the 21 bacterial isolates. Grey boxes indicate presence of genes and white boxes indicate absence of genes.

For the carotenoid biosynthesis genes, all strains possess *crtB*, while *crtA* is only present in strain WL4 and *Rhizobium* sp. strain WL3. For the *crtC* and *crtK* genes an interesting pattern is observed: *Methylobacterium* group 1 and group 2 strains contain *crtC* (with 3 exceptions, discussed below), while group 3 and 4 strains completely lack the gene. On the other hand, *crtK* is only present in *Methylobacterium* group 3 strains and missing from groups 1, 2, and 4. Similar patterns are evidenced in *pufB* which is absent from *Methylobacterium* groups 3 and 4, *Rhizobium* sp. WL3 and *Roseomonas* sp. WL45, and *pufC* missing from *Methylobacterium* group 4 as well as the novel strain WL4. The remaining genes are found ubiquitously in all strains of the analysis, with only a few exceptions (table 2). Proteorhodopsin genes were not detected in any of the strains analyzed, except for sample WL44, which is comprised of a *Roseomonas* and an Actinobacterial strain (table 1). BLAST searches showed that the identified Proteorhodopsin gene belongs to the Actinobacterial part of the mixed culture (data not shown).

The photosynthetic gene clusters were consistently found on the longer contigs of the initial assemblies of Illumina data produced by SPAdes (for the strains with fewer than 500 contigs) (table 1). The different architectures of the isolated photosynthetic gene clusters are shown in figure 6. In *Rhizobium* sp. WL3 and and *Roseomonas* sp. WL45 the photosynthetic genes form one continuous cluster, with different, however, architecture. In Methylocystaceae WL4 the genes are organized in 2 clusters, similar to all *Methylobacterium* spp. isolates. Unique to the architecture present in WL4 is the presence of *bchI* before the *crtB, crtI* genes in the first cluster. For *Methylobacterium* spp., groups 2 and 3 show similar gene synteny – though there are a few differences in gene content (table 2) – while group 1 and group 4 appear to also share the overall architecture of the photosynthetic gene clusters. For isolates WL9 and WL19 (both in group 1) the exact location of the *crtB* and *crtI* genes is not known as they appear on a short contig by themselves.

Annotating the assemblies on RAST revealed several housekeeping genes present in the vicinity of these clusters, which further supported the notion of their chromosomal placement. The completed genomes resulting from the hybrid assemblies (strain WL1, WL2, WL3, WL4, WL45) provided the final evidence of the exact placement of the photosynthetic gene clusters on the chromosomes of all our isolates. General descriptive statistics of the assembled complete genomes are given in table 3.

**Table 3:**
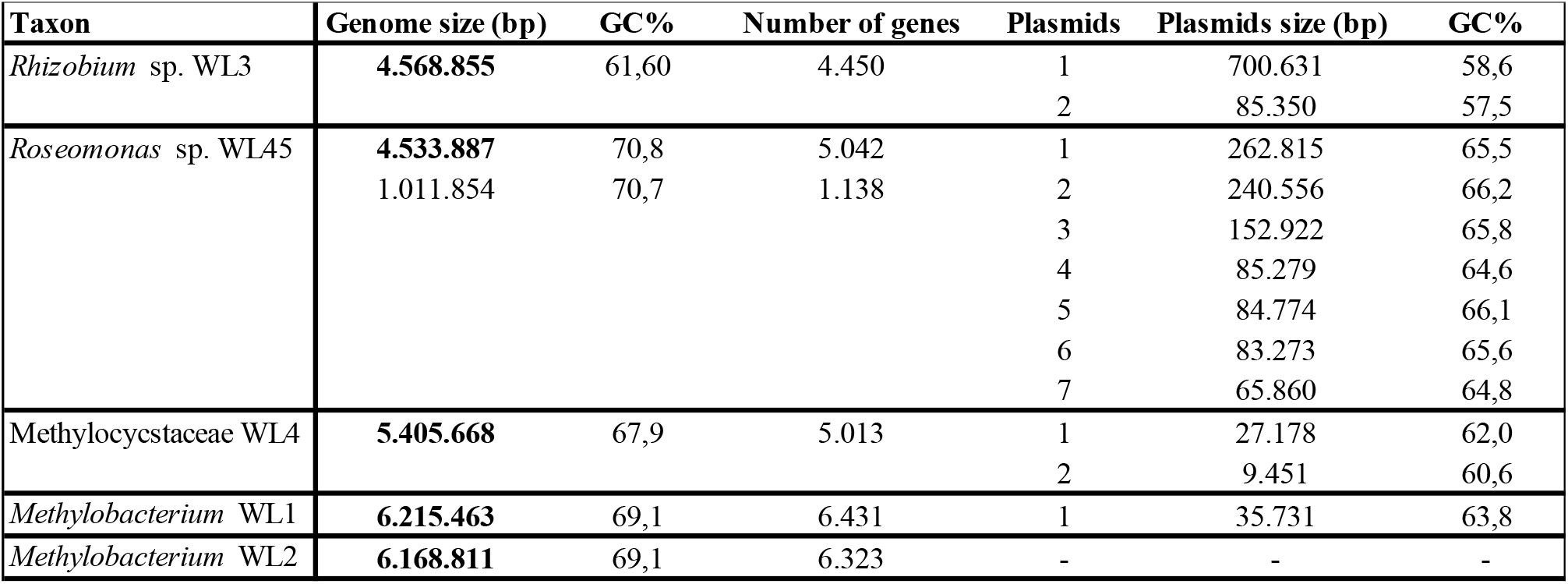
General descriptive statistics of the 5 assembled, complete AAP bacterial genomes.

The hybrid assemblies resulted in both complete genomes and plasmids. Thus, *Rhizobium* sp. WL3 contains 2 plasmids, *Roseomonas* sp. WL45 contains 2 chromosomes and 7 plasmids; the novel Mehtylocystaceae strain WL4 with 2 plasmids; *Methylobacterium* sp. WL1 with 1 plasmid; while none in *Methylobacterium* sp. WL2. Interestingly, isolates WL1 and WL2 exhibit a peculiar feature; the two isolates appear identical in the 16S phylogenetic analysis (figure 1), ANI comparison (figure 3), as well as in phylogenetic analysis of the super-matrix comprised of the 6 genes of the photosynthetic cluster (figure 5). Interestingly, the assembled genomes of WL1 and WL2 are not identical. Though close in size, WL1’s genome is slightly bigger, and it also contains 1 plasmid. Synteny between the two genomes is rather high, except for a 45kb rearrangement that is present in WL2 (Mauve alignment – data not shown). Illumina contigs show that the rearrangement in WL2 is true, based on mapping-to-reference runs. This needs to be further investigated in order to verify whether this is an artifact of the assembly process or that we truly managed to capture the beginning of divergence in these two, otherwise, identical *Methylobacterium* strains.

**Fig 5.**
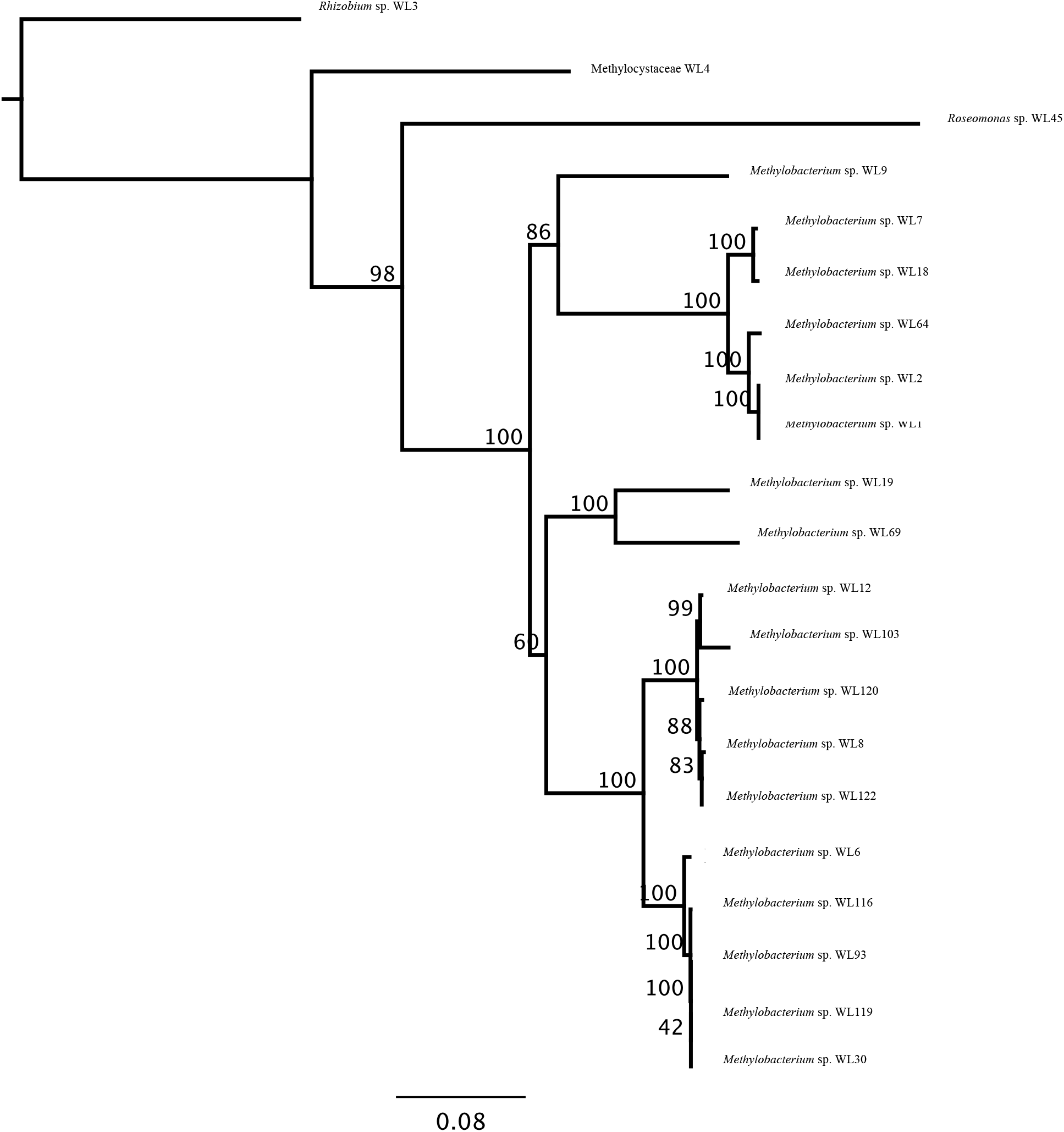
Phylogenetic tree based on the super-matrix consisting of the 16S region and the *acsF, bchL, bchY, crtB, pufL* and *pufM* genes that are present in the photosynthetic gene clusters of all 21 bacterial isolates. The legend shows substitution rates from the root of the tree and the node labels indicate bootstrap support values (%).

**Fig. 6.**
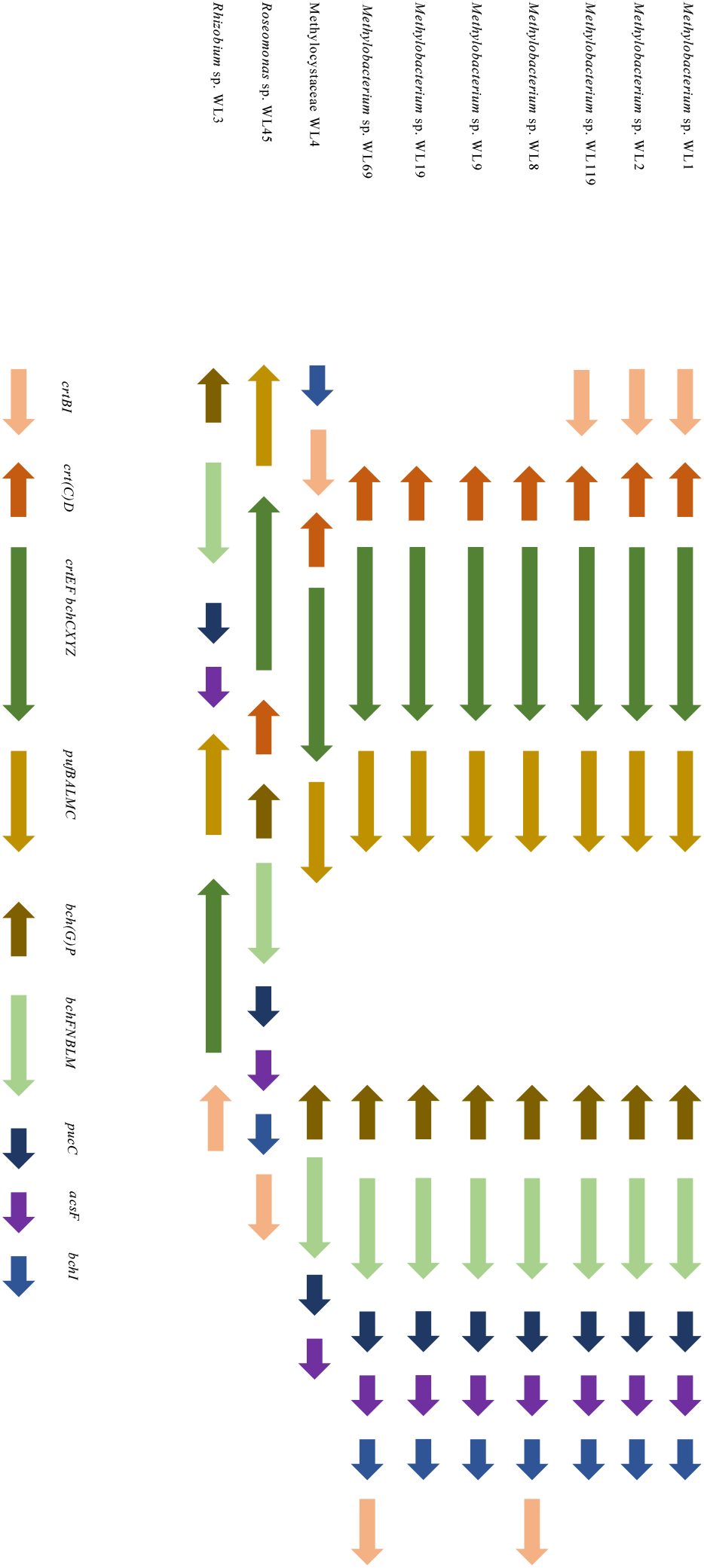
: Architecture of the different photosynthetic gene clusters. Arrows indicate the orientation of the genes. Sizes are not representatives of gene length. Genes that may be missing from some strains are denoted with parentheses on the figure’s legend.

Using *Rhizobium* sp. WL3 as outgroup and calculating substitution rates from the root of the different trees, it is observed that *Roseomonas* sp. WL45 has the most divergent genes out of the 21 isolates (table 4). This is consistent for the 16S region, the 6 genes of the photosynthetic gene clusters that were used, the super-matrix of all 7 genes, as well as the phylogenomics tree from CheckM. In *Methylobacterium* spp., which make up the majority of the isolates, it is evidenced that groups 3 and 4 have comparable rates. *Methylobacterium* spp. group 2 appear to have more divergent genes compared to the other 3 groups, with only exceptions being the genes *bchL* and *crtB*. Thus, evolution rates in the super-matrix are closer than for the individual genes. For *Methylobacterium* sp. WL103, *crtB* seems to be as divergent as that of *Roseomonas* sp. WL4, and much more different compared to the other *Methylobacterium* strains. However, the observed differences in the sequences in the selected genes of the 18 *Methylobacterium* strains disappear in the core proteome comparison (CheckM), where the margin ranges between 0.37 and 0.40 substitutions per site for all 4 groups, further supporting the clustering of these 18 isolates.

**Table 4:**
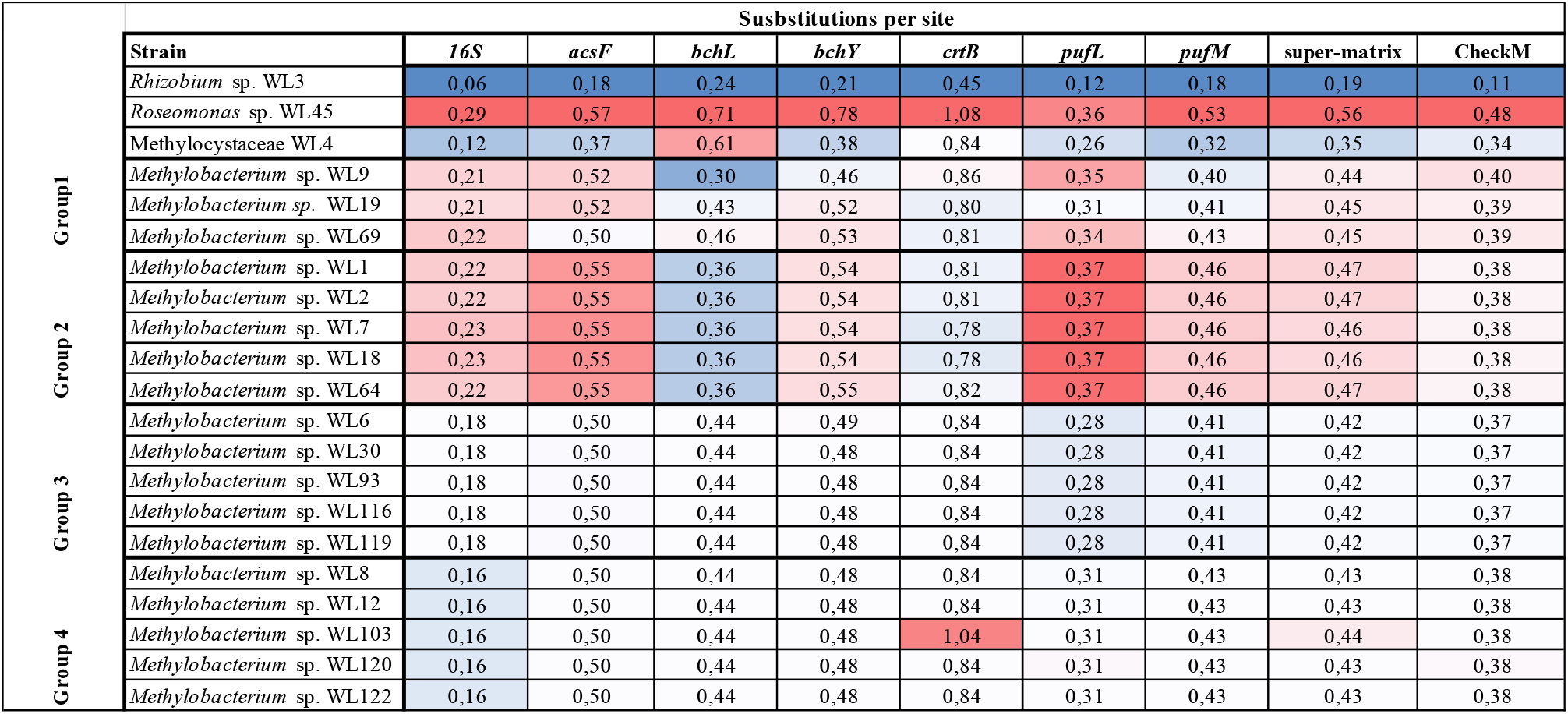
Substitutions per site in the different phylogenetic analyses. All values correspond to distances from the root, where *Rhizobium* sp. WL3 was chosen as the outgroup for all the alignments.

## Discussion

### A high throughput, multi-disciplinary approach to produce complete genomes from environmental isolates

Aiming to enhance our knowledge of the prevalence of aerobic anoxygenic phototrophic bacteria in the phyllosphere, we designed this study using wheat as the plant of choice, and employed a high-throughput, multi-disciplinary approach to identify, isolate and analyze AAP strains. The detection of bacteria bearing bacteriochlorophylls and other pigments related to photosynthesis is possible using colony infrared imaging techniques (Zeng et al. 2014). This method requires the presence of bacterial colonies growing on agar plates. It is expected that bacteria that inhabit the phyllosphere will be able to grow in such conditions in the laboratory. After identifying the “glowing” colonies, we restreaked them in new agar plates to achieve pure cultures.

From the initial 137 plates, we ended up with ~4480 bacterial colonies, more than 500 of which were probable AAP strains. Given the large number of isolates that we collected, we decided to employ a 2-step approach to reduce the total number to a more manageable figure. Thus, after macroscopic investigation of size, shape, color, and texture of the colonies, we chose 129 unique strains, for which we ran proteome profiling using MALDI-TOF MS. This approach has been shown to be able reproducibly to distinguish different taxa, and in cases where reference MALDI-TOF spectra are available in the database, be able to assign Genus and/or Species names (Madsen et al. 2015). The currently available MALDI-TOF spectra libraries lack spectra from environmental type strains (Madsen et al. 2015). Thus, we were only able to categorize the 129 isolates into 25 distinct groups. We then proceeded with whole genome sequencing on the Illumina Nextseq platform for 1 representative from each of these groups. Analyzing the draft assemblies and the 16S sequences we were able to further reduce the number of unique genomes down to 11, for which we performed Oxford Nanopore Technologies sequencing, aiming to generate complete genomes and plasmids. Following this multi-disciplinary approach that relies on classical microbiological techniques (plating, morphology etc.), biochemical analyses (MALDI-TOF MS), and two different types of high-throughput whole genome sequencing we were able to quickly, efficiently, and effectively generate draft and complete genomes of novel AAP bacteria from the wheat phyllosphere in a very short timeframe (weeks).

### Microbial rhodopsins in the wheat phyllosphere

The prevalence of AAP bacteria in the phyllosphere is not known. Studies conducted thus far relied on either metagenomes or amplicons (e.g. (Atamna-Ismaeel, Omri M. Finkel, et al. 2012; Atamna-Ismaeel, Omri Finkel, et al. 2012). Looking at bacterial rhodopsins, we identified only one in sample WL44 (~0.1% of total bacteria). The gene was shown (blast) to belong to an Actinobacterium, which however was mixed with a *Roseomonas* strain (16S analysis), which did not allow for closing its genome using ONT sequencing. Actinobacterial rhodopsins have been found in a variety of plant leave surfaces, including *Glycine max* (L.) Merr., *Oryza sativa* L., *Tamarix nilotica* (Ehrenb.) Bunge, and others (Atamna-Ismaeel, Omri M. Finkel, et al. 2012). Compared to the total number of microbial rhodopsins that were identified (156) they only constitute a minor fraction (15 sequences, ~10%). Actinobacteria, in general, grow slowly compared to other bacteria and mostly remain dormant under nutrient-limiting conditions (Barka et al. 2016), so it is possible that more rhodopsin-bearing strains are present in the wheat phyllosphere, that we were not able to detect using the infrared detection system.

In our investigation we sequenced two strains that were later revealed to belong to the anamorphic yeast *Sporobolomyces roseus* (isolates WL46 and WL96). Apart from the obvious explanation that a contamination had occurred, it is possible that these strains actually contain fungal rhodopsins, as was found earlier (Atamna-Ismaeel, Omri M. Finkel, et al. 2012). Searching for such genes under relaxed similarity criteria yielded negative results. Their draft assemblies consist of 1840 and 2078 contigs respectively, thus making them inadmissible to RAST for annotating. Moreover, a GenBank search (last accessed on 30/11/2018) using the terms “fungal rhodopsin”, “yeast rhodopsin” and *“Sporobolomyces* rhodopsin”, yielded either no results, or draft genome assemblies of yeasts with many predicted or uncharacterized opsin-type genes. We decided not to proceed with these sequences as there would be very limited biological information.

### Bacteriochlorophyll-containing AAP bacteria in the wheat phyllosphere

By investigating the presence of bacteriochlorophylls in the wheat phyllosphere, we were able to identify 21 strains harboring them, the majority of which belonged to the genus *Methylobacterium*. Members of this genus are ubiquitous in nature and have been detected in both soil and plant surfaces (Green 2006). Targeting the *bchY* (chlorophyllide reductase subunit Y) and *pufM* (reaction centre subunit M) genes in phyllosphere samples of different plants, it was shown that the Methylobacteriaceae (alpha-Proteobacteria) comprised 28% of the total *pufM* sequences from the amplicon sequencing analysis (Atamna-Ismaeel, Omri Finkel, et al. 2012). Our results are, thus, in accordance to these previous studies. As Methylobacteriaceae are indigenous inhabitants of the phyllosphere and have consistently been shown to harbor photosynthetic gene clusters, it may be that these photosystems are native parts of the Methylobacteria that inhabit this niche. Interestingly, in their study, Atamna-Ismaeel et al. had negative results from their *bchY* amplicon sequencing approach, suggesting the absence of bacteria that contain RC1 (bacteriochlorophyll-α containing reaction centers) type photosystems. Our results show that all isolates contain the complete cassette of chlorophyllide synthase subunit encoding genes (*bchB, bchC, bchF, bchG, bchI, bchL, bchM, bchN, bchP, bchX, bchY, bchZ*), as well as both the *pufL* and *pufM* genes (table 2). Testing the same primers (Yutin et al. 2009) *in silico* against our *bchY* sequences, we had positive results for all of them (data not shown). Using a whole genome sequencing approach on the positive isolates we were able to retrieve all genes contained in these photosystems and we discovered an interesting pattern: the 18 *Methylobacterium* isolates can be categorized in 4 distinct groups not only based on average nucleotide identity and whole proteome pairwise comparisons (figure 3, 4), but also from the missing genes from their photosynthetic clusters (table 2). Thus, *Methylobacterium* strains of group 1 are only missing the *crtA* (carotenoid synthase subunit A) gene, strains of group 2 are missing *crtA* and *crtK*, strains of group 3 are also missing *crtC* and *pufB*, and last strains of group 4 are additionally missing *pufC*. For the remaining 3 isolates, no patterns can be detected as we only have 1 representative from each taxon. However, all 21 isolates contain an intact chlorophyllide synthase, while different carotenoid synthase subunit genes may be randomly missing. In their Illumina assemblies, strains WL18 (*Methylobacterium* group 2), WL116 (*Methylobacterium* group 3) and WL103 (*Methylobacterium* group 4) appear to be missing more genes of the photosynthetic gene cluster compared to their closely related strains in their respective groups. This is most likely a result of poor draft genome assemblies for the 3 strains as they assembled in 1376, 1179, and 1332 contigs, respectively. Mapping their quality-controlled reads to contigs of strains of their respective groups, the missing genes had equal coverage as the rest (data not shown). Thus, we decided to report them in table 2 as present. As far as group 1 is concerned, the 3 *Methylobacterium* strains that comprise it are quite different and most likely belong to 3 distinct species, so the remaining, randomly missing *crt* genes are in this case not artifacts of a bad assembly.

Patterns of loss of the carotenoid genes have been observed in other alpha and gamma proteobacteria, like in Rhodobacterales and Sphingomonadales (Zheng et al. 2011). The absence of *crtA* results in the inability to perform the final step in the spheroidene pathway where hydroxyspheroidene is converted to hydroxyspheroidenone. Moreover, the absence of *crtC* in *Methylobacterium* spp. of groups 3 and 4 completely affects both the spheroidene and the spirilloxanthin pathways as its product is essential for the first step of both metabolic pathways starting from neurosporene and lycopene, respectively. This means that strains of these 2 groups rely on the zeaxanthin pathway to produce carotenoid pigments, and most likely nostoxanthin and erythroxanthin, which do not, however participate in the light-harvesting process, but rather have a photoprotection role (Noguchi et al. 1992; Yurkov & Csotonyi 2009).

### Phylogenetic observations and molecular evolution of the photosynthetic gene clusters

In our study we sequenced and analyzed 21 aerobic anoxygenic photosynthetic bacteria, most of which belonged to the genus *Methylobacterium*, the presence of which in the environment has already been discussed previously. *Methylobacterium* spp. are facultative methylotrophs and can utilize a variety of C1, C2, C3, and C4 compounds. Such compounds are often found in the phyllosphere due to plant metabolism (Vorholt 2012; Remus-Emsermann et al. 2014). Apart from this genus, we also identified 1 *Rhizobium* sp., 1 *Roseomonas* sp., and a strain (WL4) that belongs in the Methylocystaceae, a family of methanotroph bacteria. This is the first time that AAP bacteria from this Family have been isolated and their complete genomes assembled. Its closely related *Alsobacter* spp. have been shown to exhibit high Thallium tolerance (Bao et al. 2006), and it will be interesting to identify heavy-metal resistance genes in non-heavy-metal polluted phyllosphere samples. We propose that strain WL4 belongs in the Methylocystaceae and is in fact a novel species, and possibly a novel genus. The hybrid assembly using Illumina and Nanopore reads led to a complete genome with 2 plasmids, with the photosynthetic clusters being located on the genome. Further analysis of WL4 to verify these claims and suggest a species and/or genus name will be the focus of another investigation. However, it is worth mentioning that if WL4 contains heavy-metal resistance genes, on top of the photosynthetic gene cluster, this will have been shown for the first time and its potential to bioremediate heavy-metal contaminated areas if applied as Plant Growth Promotion and environmental remediation agent is of high value and needs to further be explored.

Phylogenetics analyses on the *Methylobacterium* groups shows that the isolates belonging in group 3 and 4 comprise a single species and possibly single strain each (figures 1, 2, 5). Group 2 *Methylobacterium* spp. possibly constitute 1 species with 2 strains (WL7 and WL18, WL1, WL2 and WL64), while group 1 isolates correspond to 3 distinct *Methylobacterium* species. This brings the total of unique isolated AAP taxa to 9, of which 5 complete genomes were assembled (WL1, WL2, WL3, WL4, and WL45). The photosynthetic gene cluster in *Rhizobium* sp. WL3 and *Roseomonas* sp. WL45 is shown as a single stretch of ~50.000 bp containing all genes. On the other hand, the novel Methylocystace strain WL4, and *Methylobacterium* sp. Strains WL1, and WL2 exhibit the bipartite architecture that has been shown elsewhere (Zheng et al. 2011). For the remaining *Methylobacterium* isolates, for which draft assemblies produced long enough contigs that would be able to encompass the photosynthetic gene cluster in one part (like WL3 and WL4), it appears that they follow a similar bipartite architecture, though it is not possible to conclude on possible differences due to the presence of some of the genes in many small contigs.

All phylogenetic trees were constructed using *Rhizboium* sp. WL3 as the outgroup. Calculating substitution rates from the root on each of the constructed trees we observed that *Roseomonas* sp. WL45 is more divergent compared to the other isolates in all analyses, including the 16S alignment as well. For the 18 *Methylobacterium* isolates, group 2 strains (WL1, WL2, WL7, WL18, WL64) evolve consistently at different rates compared to the other 3 groups. Surprisingly the trend followed by most genes in the analysis does not apply in the case of *bchL* and *crtB*. This is very interesting as all the 6 genes analyzed all belong to the same photosynthetic gene cluster. It appears that there is relatively strong selection pressure on *bchL pufL* and *pufM*, which in all cases are more similar to each other compared to *ascF, bchY* and *crtB*. This result partially comes to contrast with previously adopted methods of using *bchY* as a marker gene for RC1 clusters (e.g. Atamna-Ismaeel, Omri Finkel, et al. 2012). On the other hand, the use of *pufM* appears more sensible. It would, however, be more suitable to investigate the possibility of using other genes for metagenome amplicon sequencing approaches, such as *pufL* or *bchL*, which show a smaller degree of divergence in different genera, thus universal primers might be more successful in detecting a wide variety of AAP bacteria in environmental samples. These differences diminish when analyzing the core proteome (CheckM) of the isolates, where all *Methylobacterium* isolates fall within the range between 0.38 and 0.49 substitutions per site (table 4). Overall, it seems that the photosynthetic gene clusters were incorporated in ancerstral strains that later diverged, to give rise to the different AAP taxa. This is further corroborated by the fact that almost all phylogenetic trees (e.g. figure 5) are consistent and in agreement with both the 16S phylogenetic tree (figure 1) as well as the phylogenomic tree (figure 2). If the genes of the photosynthetic gene clusters had been transferred to these AAP several times, there would have been significant differences both in the sequence level and the phylogenetic analysis of the different genes, which would be expected to result in different topologies and diverse distances from the root, even for closely – based on 16S analysis – related strains.

### The significance of AAP bacteria in wheat phyllosphere

In this study we employed a culturomics approach to isolate for the first time a variety of AAP bacteria that are present in the wheat phyllosphere, adding to the metagenomic knowledge provided earlier from 5 other plants (Atamna-Ismaeel, Omri M. Finkel, et al. 2012; Atamna-Ismaeel, Omri Finkel, et al. 2012). As stated earlier, the phyllosphere is a rather harsh environment, with little water availability, scarce resources, very few suitable locations, extreme temperature difference between day and night, and high dosages of UV radiation (Vorholt 2012). AAP bacteria have the ability to utilize light in photophosphorylation processes, through which they produce ATP and at the same time funnel the few available nutrients in other metabolic pathways (Yurkov & Hughes 2017). This has been shown to give them an edge *in vitro* after exposure to light, compared to non-AAP bacteria (Ferrera et al. 2011). Considering that phyllosphere bacteria protect the plant from pathogenic microbes by secreting metabolites and by occupying the available niches, being able to persevere and outgrow other bacteria gives a significant edge to AAP taxa and probably points towards the suggestion of positive selection of such bacteria in their phyllosphere by the plants themselves, as they appear to be absent from the soil (Atamna-Ismaeel, Omri M. Finkel, et al. 2012). At this point, it is not clear exactly how AAP bacteria benefit the plant, especially since the phyllosphere is a complex habitat with an ever-changing, harsh environment and it required greater attention. In the past years, great interest has been shown on how to improve crop production by studying the rhizosphere. It has also been suggested that the phyllosphere communities play a significant role in plant health and growth, which ultimate leads to increased yield (Lindow & Brandl 2003). Given the current projections of world population growth and the supply of food (United Nations 2017), it becomes evident that more, in-depth studies investigating the phyllosphere of plants, and especially crops, and identifying the underlying drivers of the bacterial communities and their interactions with their plant hosts is of outmost importance. Thus, by isolating, maintaining and analyzing the genomes of phyllosphere isolates that may prove useful as plant growth promoting agents in possible field trials investigating the effect of phyllosphere inoculation to plant growth, health, and yield.

## Acknowledgements

The authors would like to thank Tue Sparholt Jorgensen for collecting the wheat sample. The authors also want to thank Tina Thane (AU), Tanja Begovic (AU) and Margit Frederiksen (NRCWE) for capable laboratory assistance. We thank Michal Koblížek for providing the initial model of the colony infrared imaging system.

## Author Contributions

AZ, YZ, LHH designed the study. YZ isolated the strains, performed colony infrared imaging, extracted DNA for Illumina sequencing. YZ and AMM performed the MALDI-TOF MS analysis. AZ extracted DNA and performed ONT Sequencing. AZ carried out the bioinformatics and phylogenetics analyses. AZ and YZ drafted the manuscript. AZ, YZ, AMM and LHH revised the manuscript for intellectual content. All authors read and approved the manuscript.

## Data availability

Complete and draft genome assemblies analyzed in this investigation are available in GenBank under the accession numbers XXXXX

